# Microfluidic trapping of vesicles reveals membrane-tension dependent FtsZ cytoskeletal re-organisation

**DOI:** 10.1101/791459

**Authors:** Kristina A. Ganzinger, Adrián Merino-Salomón, Daniela A. García-Soriano, A. Nelson Butterfield, Thomas Litschel, Frank Siedler, Petra Schwille

## Abstract

The geometry of reaction compartments can affect the outcome of chemical reactions. Synthetic biology commonly uses giant unilamellar vesicles (GUVs) to generate cell-sized, membrane-bound reaction compartments. However, these liposomes are always spherical due to surface area minimization. Here, we have developed a microfluidic chip to trap and reversibly deform GUVs into rod- or cigar-like shapes, including a constriction site in the trap mimicking the membrane furrow in cell division. When we introduce into these GUVs the bacterial tubulin homologue FtsZ, the primary protein of the bacterial Z ring, we find that FtsZ organization changes from dynamic rings to elongated filaments upon GUV deformation, and that these FtsZ filaments align preferentially with the short GUV axis, in particular at the membrane neck. In contrast, pulsing Min oscillations in GUVs remained largely unaffected. We conclude that microfluidic traps are a useful tool for deforming GUVs into non-spherical membrane shapes, akin to those seen in cell division, and for investigating the effect of confinement geometry on biochemical reactions, such as protein filament self-organization.

## Introduction

One hallmark of living entities is their ability to self-organize into and maintain complex architectures, both on the level of molecular complexes and tissues. While it has been known for decades that gradients of diffusible morphogens are crucial for the supra-cellular self-organization of tissues in development, it is now clear that similar principles operate at the intracellular level: gradients of molecules are important in cell polarity and division, and they also affect spatial dynamics of signaling.^1,2^ For example, it has been previously shown that cell geometry itself can induce a symmetry break.^3^ In eukaryotes, cell shape is controlled by both physical properties of the plasma membrane and the biochemical reactions involving components of the membrane-proximal cytoskeleton, most importantly the actin network.4 Recently, bottom-up synthetic biology has been increasingly able to reconstitute cellular phenomena in vitro, including the reconstitution of spatially organized processes.^5,6^

Reconstituting cell division in vitro represents a desirable, albeit ambitious, goal toward realizing the bottom-up construction of an artificial cell.^7–9^ In biological systems, cell division involves the segregation of chromosomes, organelles and other intracellular components, and cytokinesis, the physical splitting of the cell envelope. Bacterial cytokinesis, for example, is a complex process that requires membrane constriction and fission as well as the synthesis of new cell envelope material and separation of the peptidoglycan layer.^10^ Cell division in the vast majority of bacteria involves the GTPase protein and tubulin homologue FtsZ.10 FtsZ polymerizes into a dynamic ring-like structure at the division site, referred to as the “Z-ring”, where it is anchored to the membrane by the adaptor proteins FtsA and ZipA.^11–13^ The Z-ring serves then as a platform to recruit further divisome proteins.^7^

In addition to these membrane-transforming processes, an important aspect of cytokinesis is its spatiotemporal regulation. In order to divide at the right time and location, cells have evolved both positive and negative regulatory mechanisms to control divisome assembly.^14^ In E. coli, the MinCDE system is one of two negative regulatory systems that allow for FtsZ polymerization exclusively at the mid-cell plane. MinC is an FtsZ polymerization inhibitor that follows the pole-to-pole oscillations of the peripheral membrane-binding ATPase MinD and its ATPase activating protein MinE. These oscillations result in a gradient with highest MinC concentration at the poles and lowest at mid-cell.^15,16^

All these processes have been reconstituted in vitro, first on flat supported lipid bilayers. Here, Min D and E oscillations are visible in the form of traveling waves via an ATP-driven reaction-diffusion mechanism,^16^ and FtsZ spontaneously assembles into dynamic ring structures, in which individual FtsZ filaments undergo treadmilling to drive chiral rotations of the rings into ring-like structure.^17^ Early reconstitution experiments in spherical compartments, where only FtsZ was encapsulated in aqueous droplets in oil, showed dynamic FtsZ bundles Investigation of FtsZ inside rod-shaped droplets was also attempted, but given the microfluidics architecture only short-term observations were possible^18^. Examples of more biological meaningful compartments have been reported, such as polymersomes or giant unilamellar vesicles (GUVs).^8,19^ We and others have also reconstituted Min protein patterns in more cell-like settings, such as in PDMS microcompartments,^20–22^ in lipid droplets^23^, on the outside of lipid vesicles,^24^ and inside lipid vesicles^25^. However, bacteria are not spherical, and it is clear that cell shape has a great impact on the boundary conditions of these processes. Thus, we wondered whether FtsZ filaments would align in elongated compartments, and whether filaments would align at the neck of ‘dividing’ compartment.

Here, we present a single-layer microfluidic chip in which GUVs can be reversibly deformed into rod- or cigar-like shapes by capturing them between PDMS posts.

## Materials and Methods

### Chemicals

All reagents used in this work were from Sigma-Aldrich. Lipid compositions were prepared with DOPG (1,2-Dioleoyl-sn-glycero-3 phosphoglycerol, Avanti Polar Lipids, Inc.), EggPC (L-α-phosphatidylcholine (Egg, Chicken), Avanti Polar Lipids. Inc.), DOPC (1,2-Dioleoyl-sn-glycero-3-phosphocholine, Avanti Polar Lipids, Inc.) and DOPE-ATTO655 (ATTO-tec).

### Microfluidic device fabrication

The microfluidic vesicle trap squeezer devices were fabricated from PDMS using standard soft lithography techniques. The geometry was inspired by the traps we used in a previous study^26^, but here extended the trapping structure along the flow axis with a narrow squeezing channel containing evenly spaced indentations, and added a larger entrance funnel to increase GUV capture efficiency. We found it advantageous to design device channels with a high density of traps, which have progressively narrower funnels and squeezing channels towards the outlet.

Master moulds of 13μm height were produced on a 4 inch silicone wafer (University Wafer) using SU-8 3010 (Microchem corp.) according to the manufacturer’s data sheet and developed in PGMEA. Due to the small size of the PDMS features to be released from the mould, fluorophilic coating with Cytop was used routinely prior to the hard-baking step. In brief, 250 μL of a 1:10 dilution of Cytop CTL-809M in CTsolv.100E (Asahi Glass Co. Ltd., Japan) was dispensed on the SU-8 features and excess removed by spinning at 4000rpm for 1min. The coated wafer was then hard-baked for 30min at 180°C on a hot-plate and then allowed to cool down to room temperature slowly by turning off the heating. A 10:1 mixture of PDMS base and curing agent (Sylgart 184, Dow Corning) was homogenized and degased simultaneously for 2min using a Thinky Planetary Vacuum Mixer (ARV-310, Thinky Corp., Japan) and subsequently poured to about 4mm height onto the master in a petri-dish. After curing over-night in an oven at 75°C the PDMS was peeled off the wafer and cut to size. Fluid ports were then punched with a 3 mm and 0.75 mm diameter biopsy puncher (World-Precision-Instuments) for the reservoir and outlet, respectively. The PDMS microchannels were then sealed by plasma bonding them onto glass cover slips (24 × 32 mm, thickness 1.5, VWR) using oxygen plasma (15 s at 0.3mbar, 50% power, model ZEPTO, Diener electronic, Germany) and baking them for 15–30 min at 75 °C on a hot-plate.

### Preparation of FtsZ-containing GUVs

FtsZ samples were prepared by diluting purified FtsZ-YFP-mts27 to a final concentration of 2 μM in its buffer (25 mM Tris-HCl pH 7.5, 125 KCl, 6.25mM MgCl_2_,) and adding iodixanol (from OptiPrep™, Sigma Aldrich) at 20% was added to increase the density of the encapsulated solution, in order to improve the vesicle yield obtained.25 FtsZ samples were encapsulated in the polymeric form by adding 2 mM GTP and previously described GTP regeneration system to prolong the polymerised FtsZ lifetime28 In FtsZ experiments, the outer and inner buffer solutions were the same save an additional 180 mM glucose in the outer solution to match the osmolarity of the inner solution. (~480 mOsm/kg) (measure with Fiske Micro-Osmometer Model 210).

For FtsZ experiments, giant unilamellar vesicles (GUVs) were prepared using the water-in-oil (w/o) emulsion transfer method.29 The phospholipid solution was prepared by dissolving EggPC:DOPG, 80:20 mol % in mineral oil adding 0.02 mol % of DOPE-ATTO655 as a tracer of the lipid membrane. GUVs containing FtsZ were prepared by adding 15 μl of inner buffer into a 500 μl of phospholipid solution pipetting carefully up and down to create a homogeneous emulsion. This emulsion was deposited over a lipid monolayer previously formed for <1h between 500 μl of phospholipid solution and 500 μl of the outer buffer. The mixture was centrifuged for 7 min at 100 rcfs to deposit the GUVs at the bottom of the tube. GUVs were collected and added into the inlet reservoir of the microfluidic chip (Fig. 1).

**Fig. 1.**
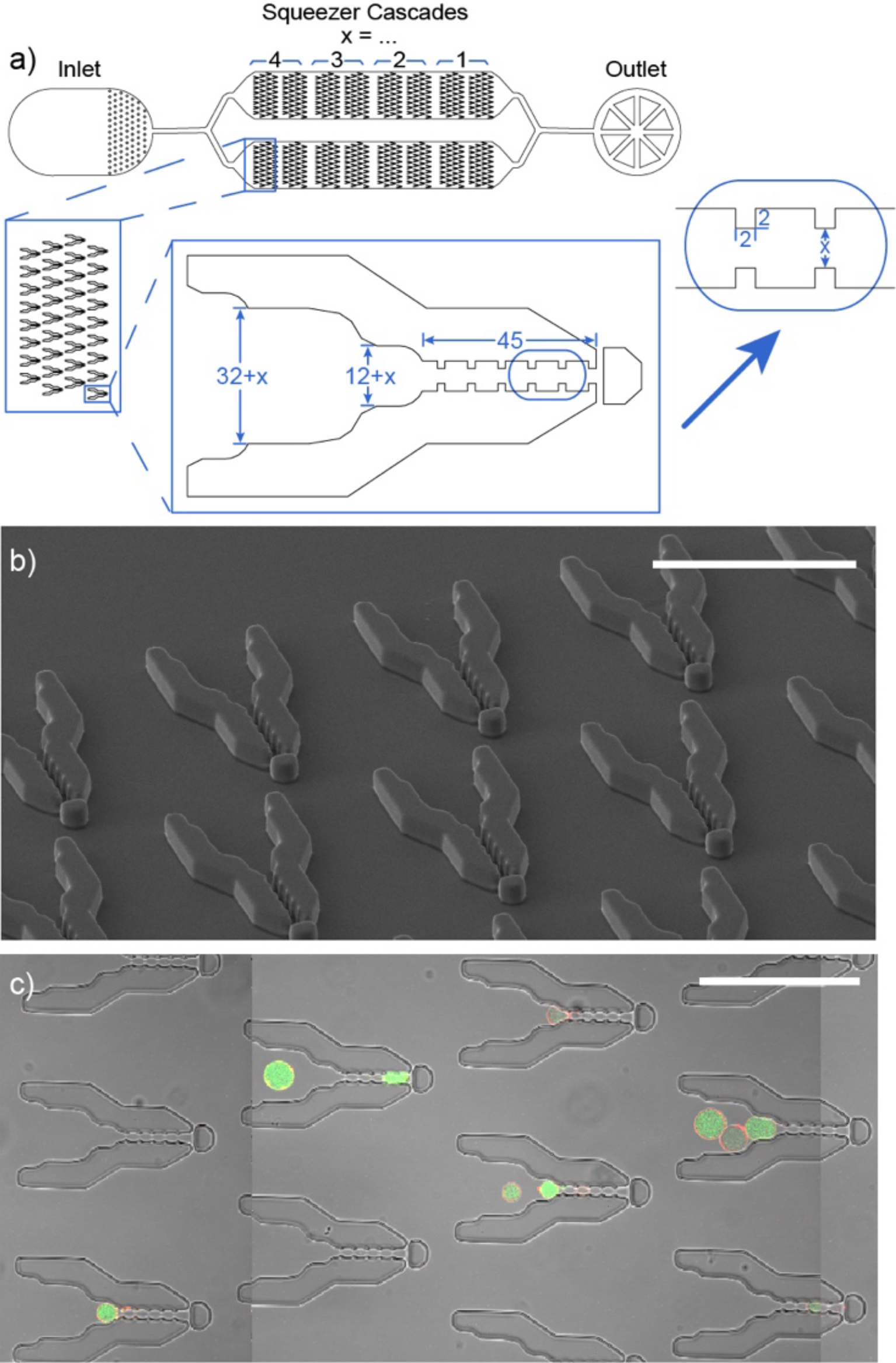
Design of microfluidic GUV traps; a) schematic depiction of chip design zooming in on the trap features. All dimensions in μm. b) SEM microscopy image of PDMS mold as a quality control step showing clean trap features including the smallest ones at the constrictions sites; c) bright field and fluorescent microscopy composite image of microfluidic chip showing the traps and trapped GUVs (containing FtsZ-YFP, green and DOPE-ATTO655 in the membrane, red). All scale bars are 100 μm.

### Preparation of Min-containing GUVs

The encapsulated Min solution was prepared by adding MinD (final concentration of 1.5 μm), eGFP-MinD (1.4 μM), MinE (3 μM) and ATP (5 μM), to a buffer containing 25 mM Tris-HCl (pH 7.5), 150 mM KCl and 5 mM MgCl2. As for the FtsZ containing vesicles, the solution contains a fraction of iodixanol (15%) to create a density gradient between GUV-inside and outside. GUVs were made using cDICE, an emulsion transfer method, first described by Abkarian et al.,30 slightly modified as described in Litschel et al..^25^ The negatively charged vesicle membrane consisted of 1-palmitoyl-2-oleoyl-glycero-3-phosphocholine (POPC) and 1,2-dielaidoyl-sn-glycero-3-phospho-(1’-rac-glycerol) (DOPG) in a ratio of 4:1.

#### Handling of microfluidic device and experimental setup

For FtsZ experiments, devices were passivated to prevent vesicle rupture upon contact with the walls by adding 20 μL of pluronic F-127 (Sigma Aldrich) at 10/50 mg/mL in phosphate buffered saline (PBS) in the inlet reservoir and centrifuging at 800 rcf for 8 min. The remaining pluronic was removed and cleaned with outer buffer. After passivation, the microfluidic devices were loaded with approximately 40 μl volume of vesicles in the inlet reservoir. A syringe joined to a pump system (neMESYS base 120 with neMESYS 290N, cetoni, Germany) and filled with 250 μl of 60% ethanol was connected to the outlet of the device, avoiding any air in between the device and the syringe. Negative flow was applied at a flow rate of approximately 5-10 μl/h to draw the vesicle solution and reagents through the fluid channels during the experiments. After 10-20 min, a high number of vesicles were collected in the microfluidic traps.. To deform the GUVs, we osmotically deflated them by replacing the buffer solution in the inlet reservoir with fresh buffer with a higher osmolarity. Some buffer exchanges with progressively higher osmolarity were sometimes necessary to induce vesicle deformation. These increments of osmotic pressure were not higher than 10-15% to avoid vesicle damage. Once the vesicles were partially deformed, the flow rate was progressively decreased to introduce the GUVs completely into the trap, deforming them slowly. To move the vesicles out of the traps and back in again, in order to see morphological differences of FtsZ in spherical or deformed GUVs, the flow rate was changed within a range of ±15-20 μL/h.

### Imaging experiments

All the experimental data were recorded with an LSM 780 confocal microscope (Carl Zeiss, Germany) equipped with a C-Apochromat, 40×/1.2 W objective. Fluorescence emission was detected by using laser excitation at 488 nm for YFP (Ftsz experiments) and eGFP (Min experiments) while 633 nm was used for Atto655. All the experiments were conducted at room temperature.

### Data analysis

All image data was displayed and contrast-adjusted (equally for all images of a given set) using ImageJ. FtsZ filaments were traced using the ImageJ plug-in RidgeDetection and the collected metadata (filament coordinates) were exported. This procedure was fully automated after setting three empirical threshold for binarisation of the image: the ‘line width’ of the filaments which was kept constant and the minimal and maximal intensity thresholds, which were chosen by visual inspection and comparison of the detected filaments and the original image. Custom-written MATLAB code was used to extract the angles between filaments and the long GUV axis as well as filament length distributions, separating the ‘neck’ region from the remainder of the GUV by a manually selected ROI and to generate the plots shown in the figures. For the quantification of the aspect ratio, the lengths of the GUVs’ x and y axis was extracted using ImageJ while MATLAB was used for the statistical analysis.

## Results and discussion

### Design and optimization of the microfluidic device

The microfluidic design consists of 2 channels with 280 traps and was based on previous design for capturing GUVs.^26,31^ After several design iterations, we found that the design shown in Fig. 1 – asymmetric traps with a large funnel – were optimal for GUV capture (see also Figure S1a, design FS814). Adjacent columns of traps are vertically offset from one another to maximise GUV capture. The spacing of the traps is reduced in four steps in the direction of the chip outlet, so GUVs of different sizes can be efficiently captured. Since we are interested in reconstituting elements of a synthetic cell division machinery in these elongated GUVs, we also designed the traps to mimic the ‘neck’ of the division site (Fig. 1a). Each trap has multiple indentations in order for GUVs to sit more stably in the trap, and a ‘stopper’ to prevent GUVs from escaping. Since the constriction features are small and have a high aspect ratio (2×2×13 μm), we made sure that our final traps faithfully replicated our design by imaging both the wafer using Laser Scanning based Profilometry (Fig. S1b) and the PMDS mold using scanning electron microscopy (Fig. 1b).

### Deforming GUVs containing membrane-bound FtsZ filaments results in their reorganisation on the GUV membrane

In order to test how encapsulated FtsZ filaments respond to a different compartment geometry, we made GUVs containing purified FtsZ in polymerizing conditions using an emulsion transfer method.^17,32,33^ After pipetting a suspension of GUVs into the inlet reservoir, we used a syringe pump to apply flow from the outlet to suck GUVs into the channels after pipetting a solution of GUVs into the inlet reservoir. We were able to capture GUVs at a constant flow rate of 5 μl/h and keep them trapped for minutes (Fig. 1c). Some of the captured GUVs already started to deform at these flow rates, going from a spherical shape to a slightly elongated shape (Fig. 1c, white arrow). However, in most cases the GUVs were not fully deformed. While we succeeded in ‘squeezing’ the GUVs fully into the narrow funnel of the traps by using up to 50× the flow rate used for loading the traps, the trapped GUVs would soon after be either pushed out of the traps, or revert to a more spherical shape as soon as the flow rate was reduced. However, when we applied a mild osmotic deflation (14-23% increase in buffer osmolarity), GUV deformation was greatly facilitated and, depending on their size, some of them remained stable in a rod-like shape (change of mean aspect ratio from ~1 for spherical GUVs to 6.7±1.9, Fig. 2a-c and Fig. S2a). We were thus able to create GUVs with a height of 13 μm (height of trap channel), width of 7 μm (or 5 μm at the constriction sites) and arbitrary lengths, up to ~50μm (Fig. S2b). This allowed us to trap and deform GUVs. We expect the osmotic deflation to result in slight volume loss of the GUV, and assume that the concomitant increase in membrane area allowed the GUV to be stable in a non-spherical shape.

Although we could not detect a volume change by confocal microscopy, this seems to be the most likely explanation for the observation of stable, elongated GUVs inside our microfluidic traps. While FtsZ intensity did not change significantly upon osmotic deflation and GUV deformation, the shape change caused the FtsZ filaments to align preferentially perpendicular to the GUV long axis, particularly at the GUV neck (Fig. 2d). Thus, imposing a rod-like geometry with a central membrane neck seems to be sufficient to align FtsZ filaments at the neck location, mimicking FtsZ assembly into the Z-ring at the division site in live bacteria. To investigate the effect of GUV geometry on FtsZ filaments organisation in the absence of an osmotic deflation, we tested whether it was possible to bring the elongated GUVs back into a spherical shape by reversing the direction of flow to flush the GUVs out of the traps. Strikingly, a minor change from negative to positive pressure push most of the GUVs back outside the narrow compartment, where they reassumed a spherical shape (Fig. 3a, b-c). Thus, the initial deformation was fully reversible. In addition, the sequence of deformation and relaxation into spherical shape could be repeated multiple times (Fig. 3a, d-e). These subsequent deformation cycles simply required changes to flow direction and velocity, in stark contrast with the initial loading and deformation step. Once the GUVs gained back their spherical shape, we also observed a change in FtsZ filaments organization (Fig. 3 b-c). FtsZ filaments reorganised from elongated and curved filaments into dynamic, ring-like structures (Fig. 3c,e – FtsZ channel), resulting in, on average, shorter filament lengths (Fig. 3f). This is likely caused by the low membrane tension and release of extra membrane area, which is expected if an elongated GUV reassumes a spherical shape at constant volume. Interestingly, the reorganisation into the dynamic rings in spherical GUVs also resulted in the formation of membrane protrusions, consistent with this idea (Fig. 3c,e – membrane channel, Fig. S3). Just like the overall GUV shape, this FtsZ reorganisation was fully reversible: if a GUV with ring-like FtsZ filaments assumed a rod-like shape again, the long FtsZ filaments reformed (Fig. 3d), and if the GUV was released again, the rings reformed from the filaments (Fig. 3e). The long FtsZ filaments were predominantly aligned perpendicular to the GUV long axis (Fig. 3g). These experiments showed that in the case of FtsZ, geometry plays an important role in filament organisation, with a break in symmetry guiding FtsZ filaments to align along the short axis, in the same fashion as in vivo. FtsZ filaments are thus likely able to sense geometry, as well as membrane tension, as shown by the (reversible) transition from FtsZ filaments to rings on the membrane surface upon reduction of membrane tension and release of extra membrane area.

**Fig. 2.**
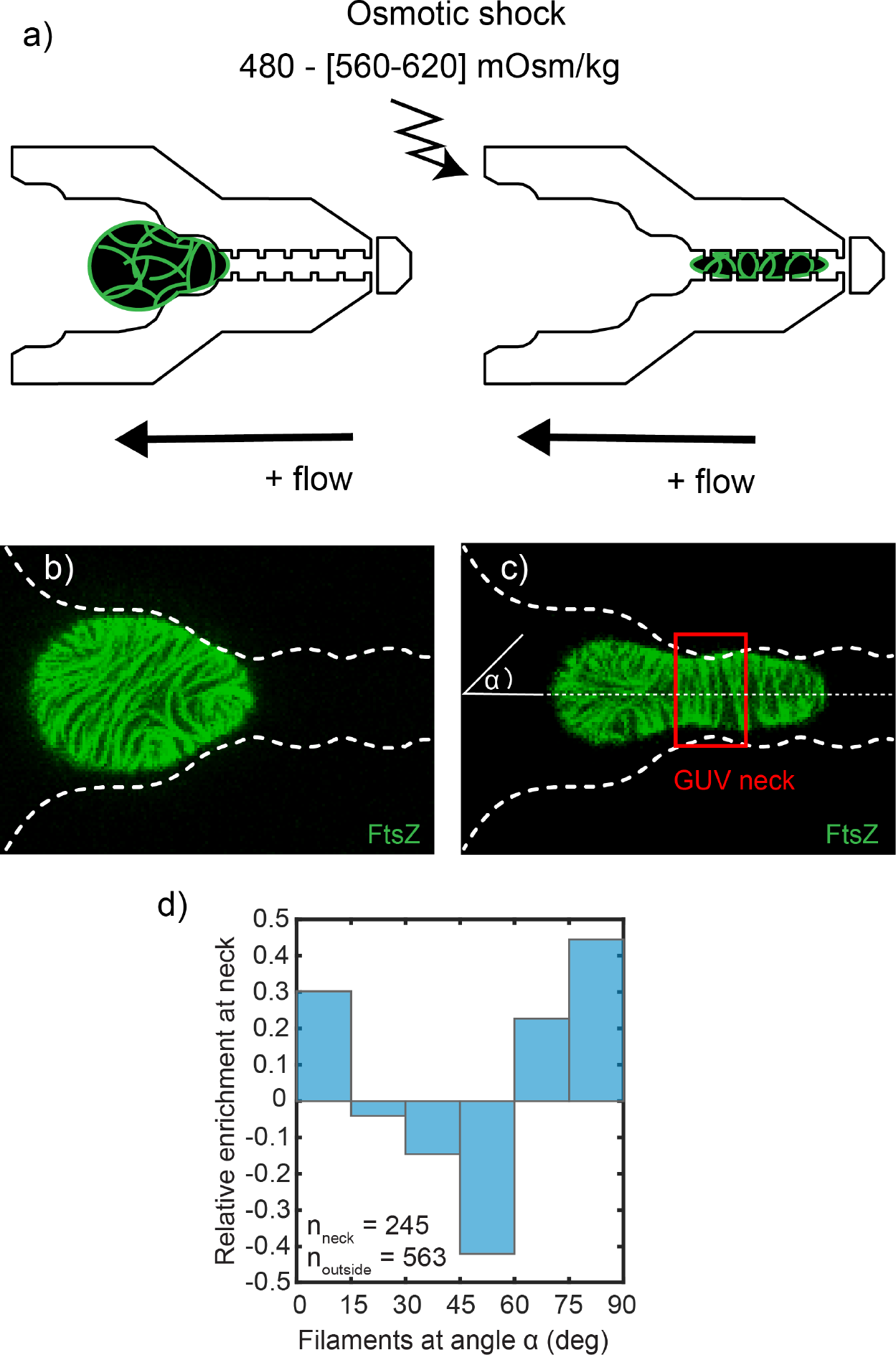
Elongation and indentation of GUVs containing FtsZ filaments leads to filament alignment orthogonal to the long axis at the GUV neck; a) schematic depiction of experiment; b), c) confocal images of the equatorial plane of trapped GUVs, FtsZ is shown in green and sub-stoichiometrically labelled with YFP. b) before and c) after osmotic deflation and flow induced deformation; d) relative enrichment of filament angles α at the GUV neck compared to other parts of the GUV. n given is the number of filaments analysed; n(GUVs) = 12 from 3 independent experiments.

**Fig. 3.**
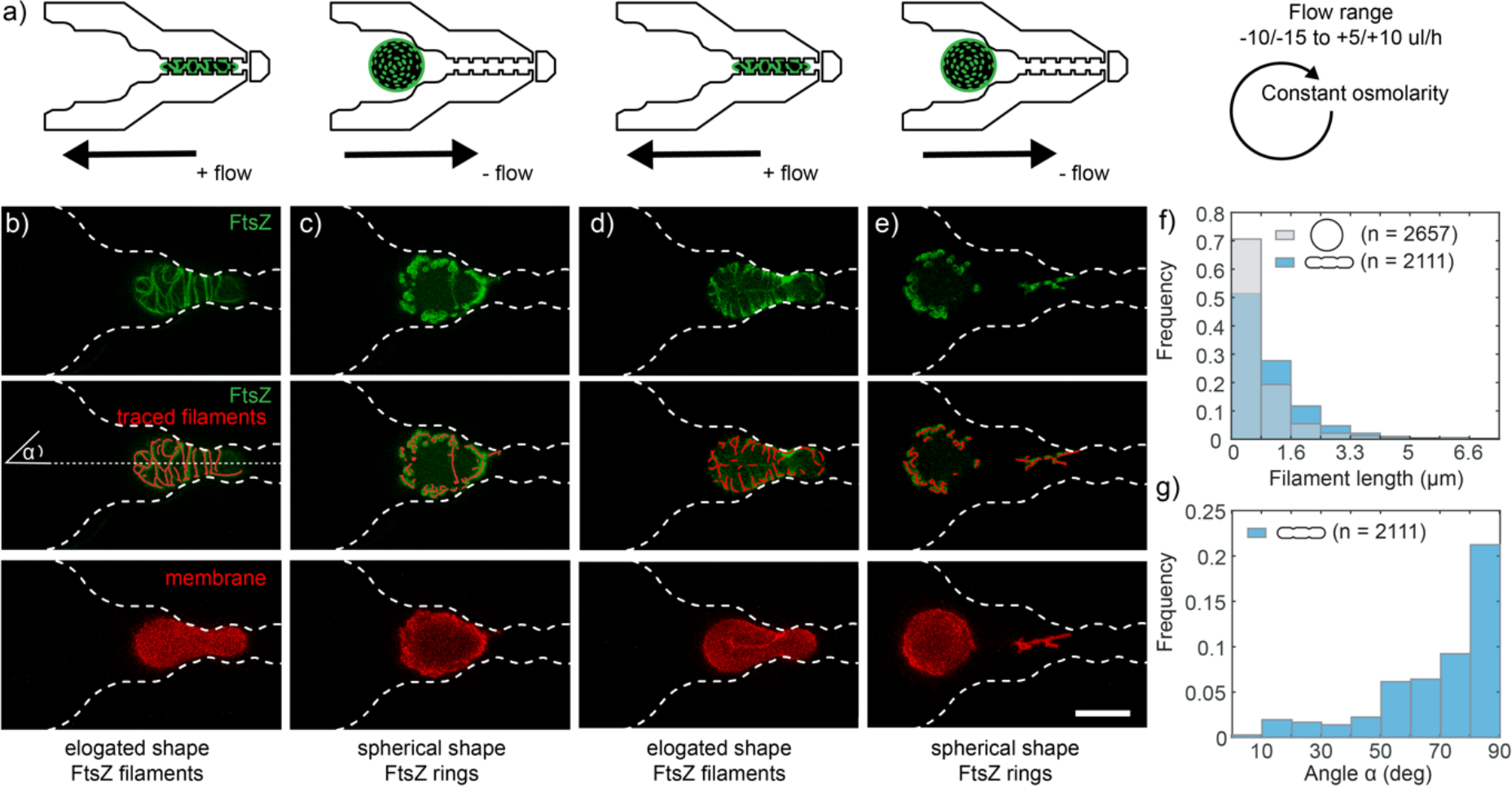
Reversible elongation and relaxation of FtsZ GUVs back into spherical shapes leads to reversible transitions between FtsZ rings and long filaments; a) schematic depiction of experiment; b) confocal images of the equatorial plane of a trapped and elongated GUV showing FtsZ filaments (top), traced filaments (middle) and GUV membrane (bottom); FtsZ is shown in green and sub-stoichiometrically labelled with YFP. Trap outline is marked (white dotted line). c), d), e) transition of the same GUV between spherical c),e) and elongated d) shapes purely by manipulation of flow rates and direction. f) Distribution of filament lengths for elongated and spherical GUVs, illustrating the filament to ‘ring’ transition. n given is the number of filaments analysed; g) distribution of filament angles α for elongated GUVs; n(GUVs) = 12 from 3 independent experiments. The scale bar is 10 μm.

### Pulsing Min oscillations in GUVs are not affected by GUV deformation

Recently our group reported the encapsulation of the Min protein system inside GUVs, thus allowing us to investigate the effect of geometry on this protein system within deformable vesicle compartments.^25^ The aspect of geometry sensing of the Min system has been studied before by restricting the reaction space to patches of SLBs^34^ or in membrane-lined microfluidic chambers.^20,22^ However, these methods are limited to non-deformable compartments, precluding changes to compartment geometry during oscillations and falling short of the possibility to reconstitute the Min oscillations on freestanding membranes. Similar to our experiments with FtsZ-containing GUVs, we encapsulated the proteins MinD and MinE into GUVs and were able to trap these within the microfluidic traps. By using again a combination of osmotic deflation and flow rate control, we were able to reversibly deform the GUVs into elongated shapes (Fig. 4a). In contrast to the FtsZ experiment, we osmotically deflated the GUVs not only for the first deformation event, but also at every subsequent time when the GUVs were squeezed into the microfluidic traps. In an extreme case, a large vesicle filled the entire length of the trap channel but still had an undeformed region (Fig. S4). Min-containing GUVs can exhibit different oscillation modes.25 We examined the effect of deformation on the pulsing mode oscillation, which represents the most common oscillation type. In these vesicles, all encapsulated proteins oscillate in synchrony between two states: MinD is either freely diffusing in the vesicle lumen (Fig. 4a, panels 1, 3 and 5) or it is bound the inner membrane surface (Fig. 4a, panels 2,4 and 6). Interestingly, when we followed the oscillations by measuring the fluorescence intensity of the vesicle lumen over time vesicle elongation did not result in any qualitative changes in oscillation dynamics for this oscillation type (Fig. 4b). In fact, the frequency and amplitude of the oscillations was remarkably constant (Fig. 4c, Fig. S4, S5). Transitions to a different oscillation mode were also not observed. Since we calculated the amplitude and frequency by averaging over the entire GUV, we wondered whether this analysis could mask spatial heterogeneity in the elongated GUVs. For example, there could be a delay in oscillation onset (a phase shift) for front compared to the back end of the GUV. Therefore, we analysed the oscillation for three regions of interest along the long axis of a deformed GUV (Fig. S4). We found that both amplitude phase were similar along the long axis of deformed GUVs (Fig. 4e). Only for the case of the largest GUV trapped and deformed (diameter ~30 μm, Fig. S4), we observed a small phase shift between the front and the back end of the GUV (~4% of period, Fig. 4e and S4).

**Fig. 4.**
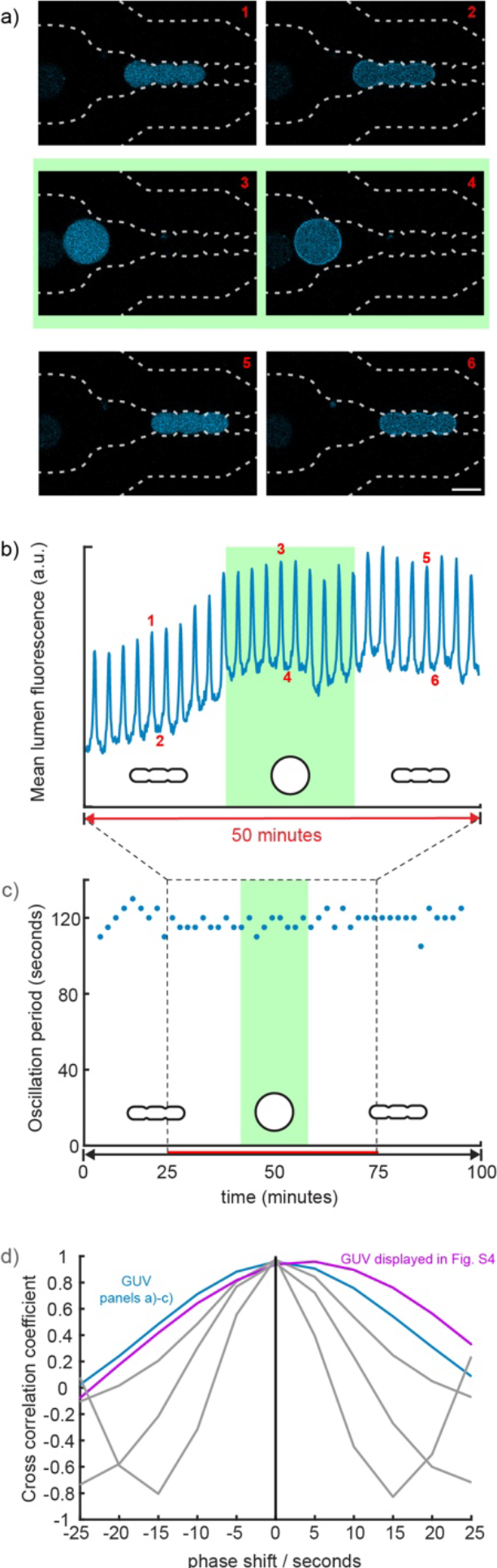
Trapping and elongation of GUVs containing the oscillating Min protein system. a) Time series of confocal images showing trapped GUVs; MinD is labelled with eGFP (cyan), trap outline is marked (white dotted line). The 6 snapshots show an elongated GUV (1-2), the same GUV in a relaxed spherical state after reversing flow (3-4), and finally again in an elongated state (5-6). For each geometry, the two states of the protein oscillation are shown: odd-numbered images show the bulk of the MinD in the vesicle lumen and even-numbered images show membrane-bound MinD. Scale bar is 10 μm. b) Average fluorescence intensity of vesicle lumen over time. Green shaded area indicates time during which the GUV is not deformed. c) Period (time between two intensity peaks) of Min oscillations over more than 50 cycles. d) Cross-correlation of the signals from the ROIs drawn at opposite ends of squeezed GUVs for different lag times (phase shifts; see also Fig S4 and S5).

## Conclusions & discussion

We have designed microfluidic traps for deforming GUVs with constriction sites, mimicking the indentation of a division furrow. We were able to create GUVs with a height of 13 μm, width of 7 μm (or 5 μm at the constriction sites) and arbitrary lengths. This means that we were able to change the mean aspect ratio from ~1 for spherical GUVs to 6.7±1.9 for trapped and deformed GUVs, close to that expected for rod-like shapes. Parallel to this work, Fanalista et al. have developed microfluidic traps to deform single- and double-emulsion droplets to spheroid-like shapes, covering a wide range of the morphological space as defined by the droplets’ aspect ratio and volume.^35^ In the future, we would therefore expect that a similar parameter space could be covered for deforming GUVs by combining a multi-height chip similar to theirs with our design and approach, going down to channel heights similar to the length of the narrowest constriction sites (<5 μm). As a complimentary approach to microfluidic traps, our lab has also developed microstructures formed by a 3D-printed protein hydrogel. Shape changes of trapped GUVs are then reversibly induced upon pH-induced swelling of the hydrogel (paper submitted). In this work, however, GUVs were deformed into quadratic, triangular or disc-like shapes with an aspect ratio of ~2.

As the occupancy of our chip’s 280 traps can be adjusted by GUV concentration and loading times, it is possible, in principle, to deform many GUVs in parallel. In practice, however, we have always focussed our observation on a single GUV at a time, as doing so allowed us to resolve the GUV’s response to the shape changes at maximal resolution. In future experiments, data from many GUVs could be obtained in parallel if the read out does not require high-resolution confocal imaging. However, the high number of traps was still an advantage for our single-GUV imaging experiments, allowing us to easily select a GUV with the required properties (e.g. encapsulated FtsZ) from heterogeneous GUV population. GUVs trapped in our microfluidic chip could be observed for long times (100 minutes). We showed that, after an initial osmotic deflation, deformation was fully reversible, making it possible to study whether shape-change induced effects were also reversible.

We found that this was the case for the shape-induced reorganisation of encapsulated FtsZ filaments. Upon the first, osmotic-deflation induced, deformation of GUVs that contained elongated FtsZ filaments, FtsZ filaments merely reorganised in their spatial orientation. When these rod-shaped GUVs were released from the traps, the FtsZ filaments reorganised into ring-like structures while the GUVs reassumed a spherical shape. This can be most likely explained by the increase in apparent membrane area as the GUVs relaxed to a shape with a lower surface/volume ratio. Controlled deformation of GUVs in microfluidic traps is therefor also a way to manipulate membrane area and tension, and to study its effect on membrane-protein interactions and membrane-bound protein networks.

Analogously to the FtsZ experiments, we examined the effect of GUV deformation on an encapsulated reaction-diffusion system (the Min protein system). In particular, we found that the pulsing mode oscillation, which represents the most common oscillation type, was unaffected by GUV aspect ratios. We note that in our experiments, the GUVs are osmotically deflated and we therefore expect to be in a regime of low membrane tension. Hence, the increase in surface-to-volume ratio upon GUV deformation/elongation is due to an increase in apparent membrane area by flattening out membrane undulations. This means that the total membrane area is conserved and that the actual membrane area/volume ratio remains constant: instead, the overall geometry and local membrane architecture is changed. It is interesting that changing the aspect ratio alone is not enough to influence the oscillations as one may have expected (as judged from their frequency/amplitude or their type, e.g. pulsing vs. pole-to-pole oscillation).

## Outlook

Overall, we present this device as a platform for studying the effect of compartment geometry on processes reconstituted in and on the membrane of GUVs or minimal cells. Beyond GUVs, our traps could also be used to deform droplets or vesicle-like membranes derived from cells, such as spheroblasts36 and giant plasma membrane-derived vesicles.^37,38^ In the present study, we have investigated the effect of GUV shape on membrane-bound FtsZ filaments and the Min reaction-diffusion system. In the future, it would be interesting to study the shape-dependent effects on other reaction-diffusion and cell-polarity inducing systems, such as Cdc42.^39^ Equally, the effects on other cell division systems than FtsZ, e.g. MreB, or cellular signalling, could be elucidated.^40^ With the increasing capacity to create artificial, minimal cells, e.g. by encapsulation of cell-free expression systems,^41^ a device such as the one presented herein could be used for studying the effect of compartment geometry on more and more complex molecular networks.

## Acknowledgements

The authors are grateful to Germán Rivas (CSIC, Madrid) for helpful discussions. We acknowledge the MPI-B Biochemistry Core Facility for assistance in protein purification. K.A.G. has received funding from the European Union’s Horizon 2020 research and innovation programme under the Marie Skłodowska-Curie grant agreement No. 703132. D.A.G-S. and P.S. acknowledge financial support from the DFG through the Graduate School of Quantitative Biosciences Munich (QBM) This work is part of the MaxSynBio consortium, which is jointly funded by the Federal Ministry of Education and Research of Germany (BMBF) and the Max Planck Society (MPG).

## Supplementary Material

**Figure S1.**
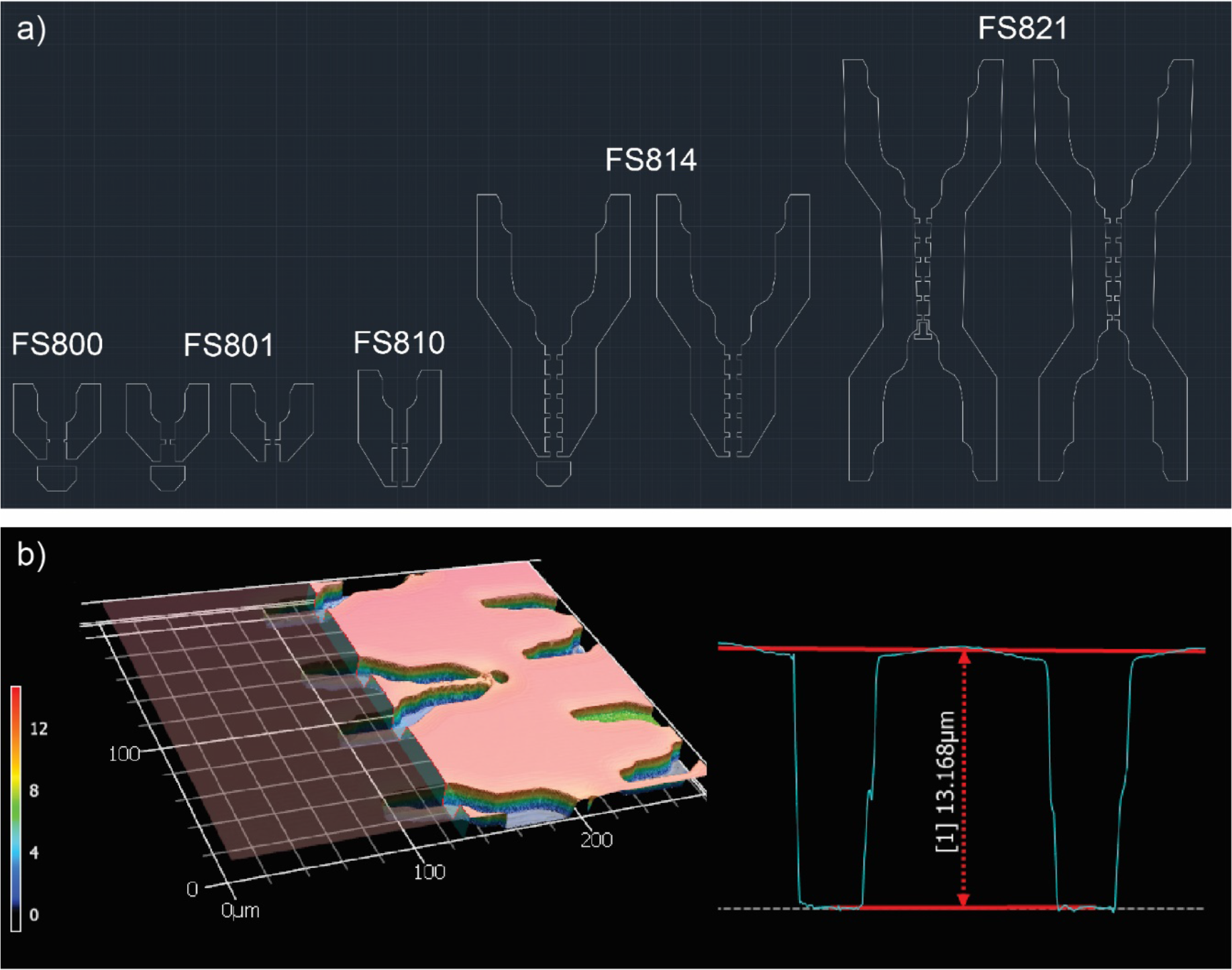
Design of microfluidic GUV traps; a) different iterations of chip design; b) profilometry microscopy image of SU8 features on wafer for quality control.

**Figure S2.**
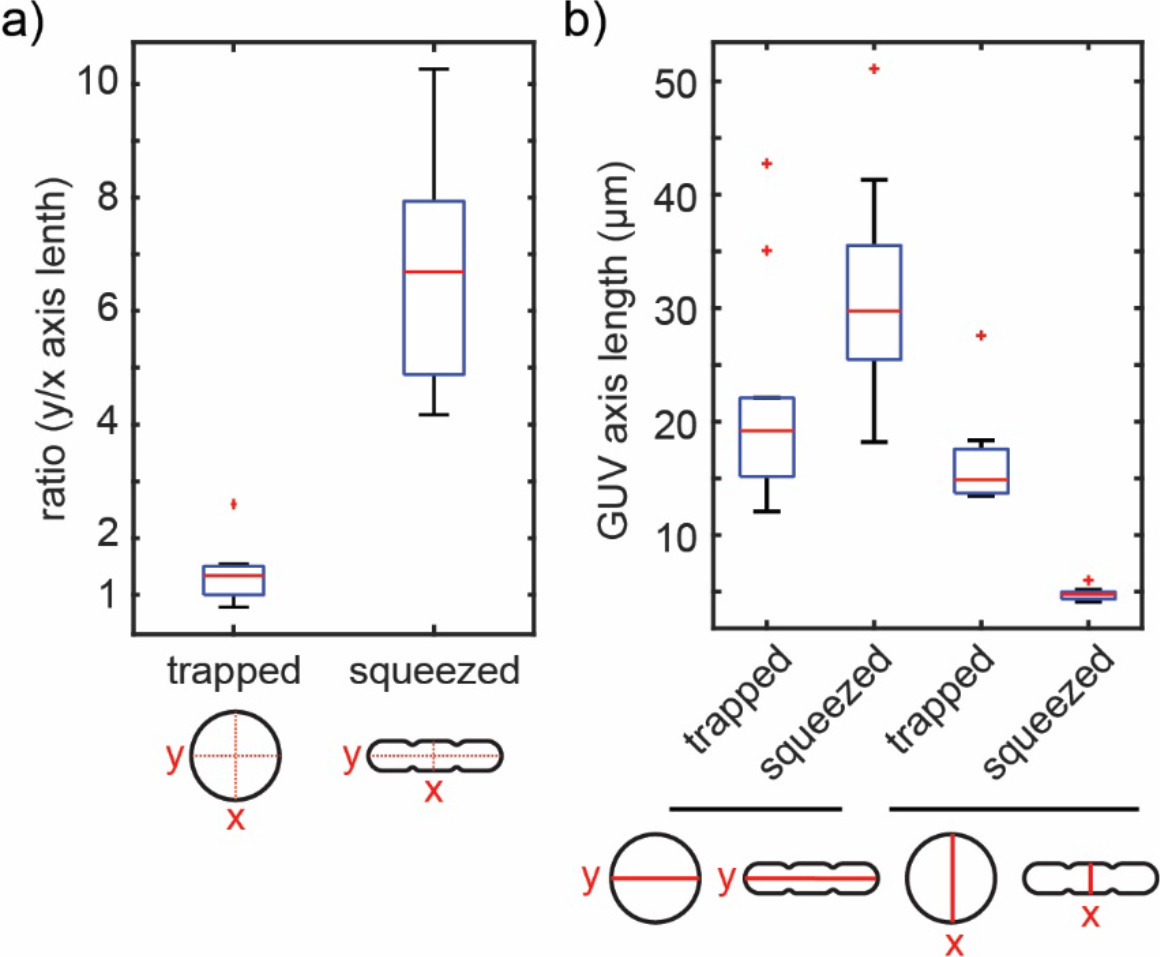
Determination of GUV aspect ratios after trapping and after squeezing; a) y/x axis length ratio; b) x and y axis length (input for the y/x ratio plotted in a)) were manually determined in ImageJ at GUV mid plane. Box plot denotes median in red, interquartile range as blue box, the 2.7σ (99.3%) confidence interval as whiskers and outliers as red dots. n(GUVs) = 10.

**Figure S3.**
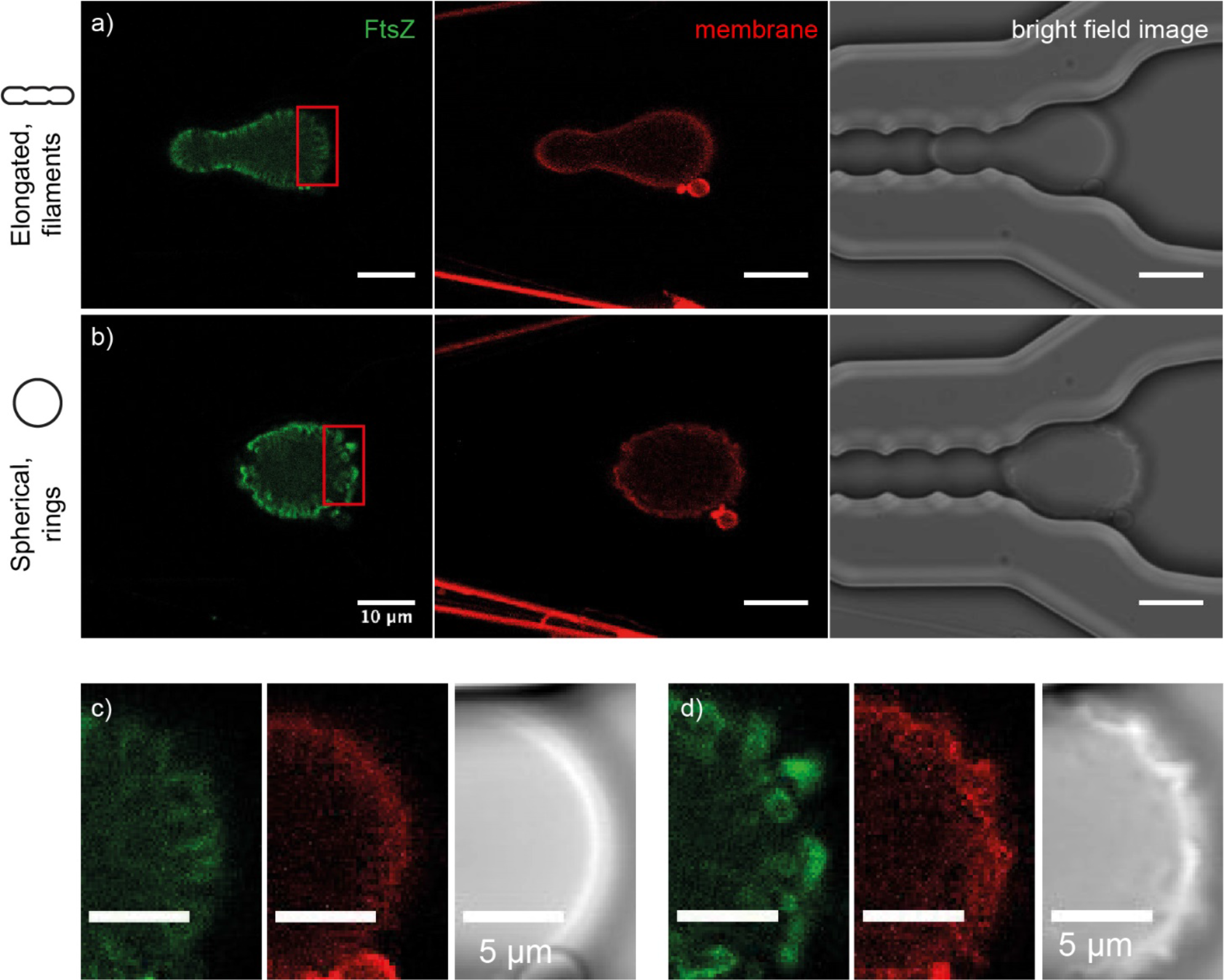
FtsZ reorganisation into dynamic rings upon relaxation back into spherical shape results in membrane protrusions. a) confocal and corresponding bright field microscopy images of elongated GUV (membrane shown in red, DOPE-ATTO655) containing FtsZ filaments (green); b) the same GUV imaged after release from the trap, having assumed a spherical shape again; c),d) close-up images of the regions marked with a white rectangle in a) and b). Note that the membrane shows protrusions after FtsZ filaments have rearranged into dynamic rings d) that are absent in c).

**Figure S4.**
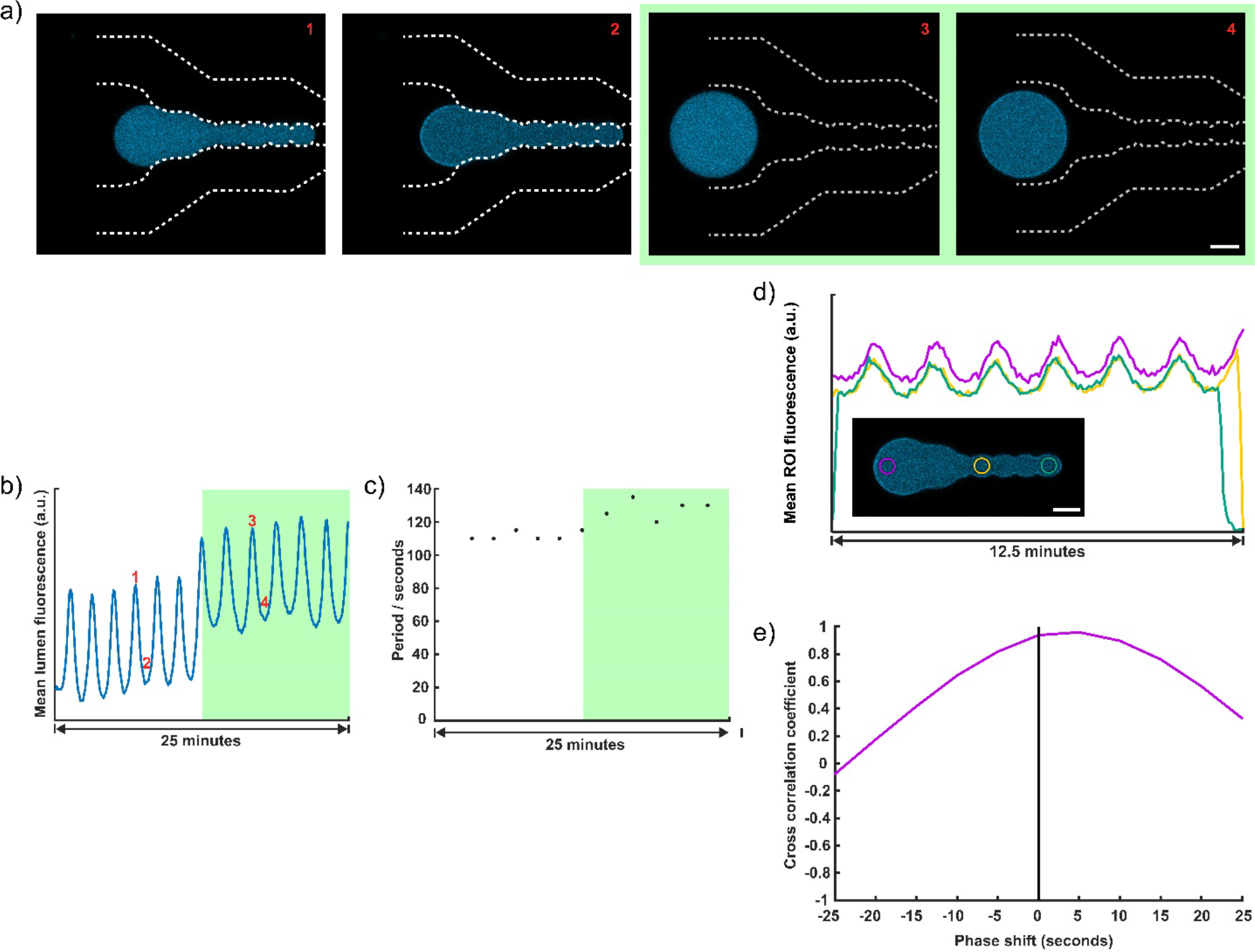
Spatial homogeneity of Min oscillations in the elongated GUVs. a) Time series of confocal images showing trapped GUVs; MinD is labelled with eGFP (cyan), trap outline is marked (white dotted line). The 4 snapshots show an elongated GUV (1-2) and the same GUV in a relaxed spherical state after reversing flow (3-4). For each geometry, the two states of the protein oscillation are shown: odd-numbered images show the bulk of the MinD in the vesicle lumen and even-numbered images show membrane-bound MinD. Scale bar is 10 μm. b) Average fluorescence intensity of vesicle lumen over time. Green shaded area indicates time during which the GUV is not deformed. c) Period (time between two intensity peaks) of Min oscillations over more than 12 cycles. d) Traces of fluorescence intensity over time of three regions of interest (ROI) in vesicle lumen of a GUV spanning four constrictions sites (insert, MinD-eGFP shown in cyan) and with a >five-fold change in aspect ratio. The plotted signals correspond to the ROIs marked in violet, yellow and green, respectively, in the microscopy image. The traces show almost perfect agreement in amplitude and frequency of the Min oscillation. e) Cross-correlation of the signals from the three ROIs shown in d) reveals that the phase of the oscillation is shifted by ~5 seconds for ROIs at opposite GUV ends (violet, green).

**Figure S5.**
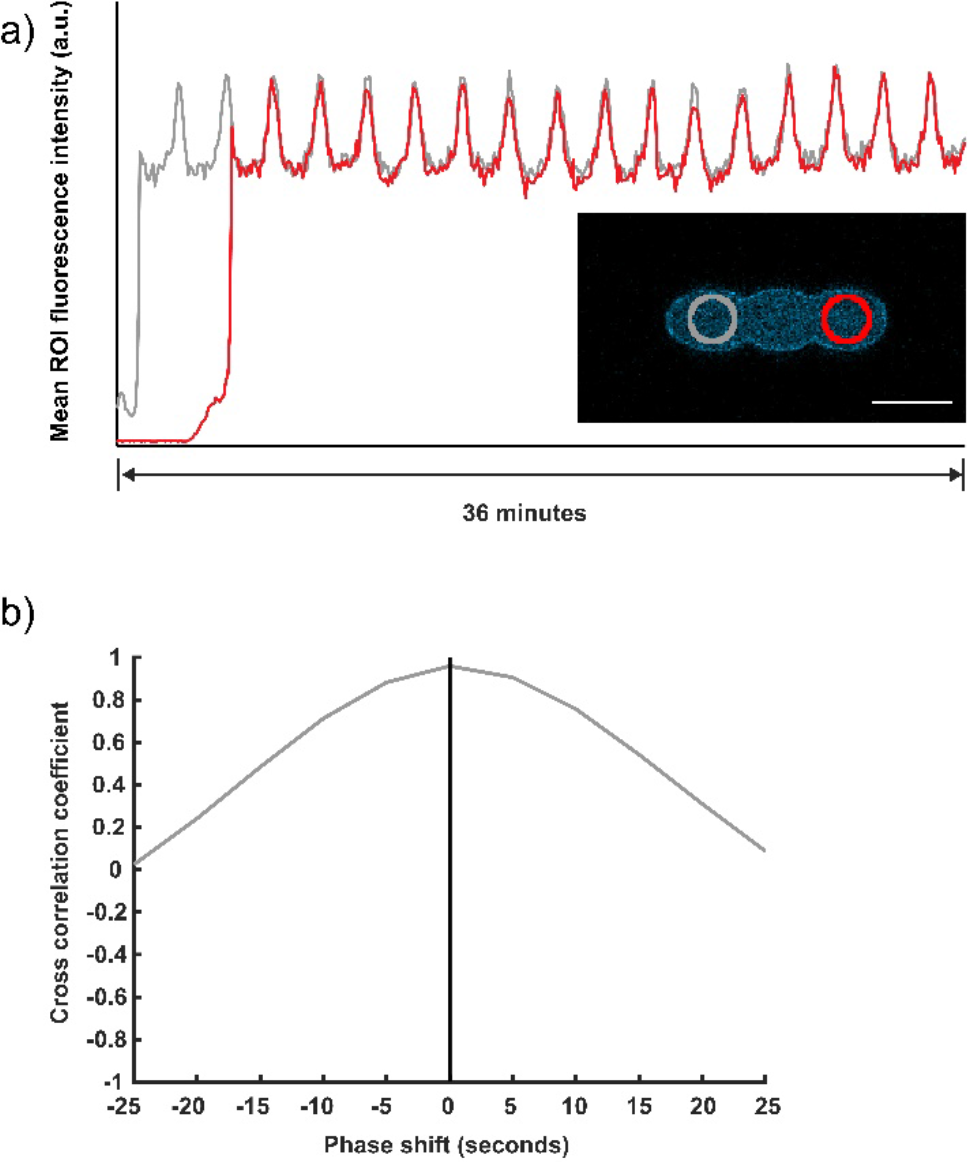
Spatial homogeneity of Min oscillations in the elongated GUVs. a) Traces of fluorescence intensity over time of two regions of interest (ROI) in vesicle lumen of the GUV shown in Fig. 4 spanning two constrictions sites (insert, MinD-eGFP shown in cyan). The plotted signals correspond to the ROIs marked in red and grey, respectively, in the microscopy image. The traces show almost perfect agreement in amplitude, frequency and phase of the Min oscillation. b) Cross-correlation of the signals from the ROIs shown in a) reveals that the phase of the oscillation is similar for both ROIs.

